# Long-Term Oral Tamoxifen Administration Decreases Brain Derived Neurotrophic Factor in the Hippocampus of Female Long-Evans Rats

**DOI:** 10.1101/2023.04.26.538155

**Authors:** Laura E. Been, Amanda R. Halliday, Sarah M. Blossom, Elena M. Bien, Anya G. Bernhard, Grayson E. Roth, Karina I. Domenech Rosario, Karlie B. Pollock, Petra E. Abramenko, Leily M. Behbehani, Gabriel J. Pascal, Mary Ellen Kelly

## Abstract

Tamoxifen is a selective estrogen receptor modulator (SERM) that is commonly used as an adjuvant drug therapy for estrogen receptor-positive breast cancers. While this drug is effective at reducing the rate of cancer recurrence, many patients report unwanted cognitive and affective side effects such as brain fog, confusion, memory impairment, anxiety, and depression. Despite this, the impacts of chronic tamoxifen exposure on the brain are poorly understood, and rodent models of tamoxifen exposure do not replicate the chronic oral administration seen in patients. We therefore used long-term *ad lib* consumption of medicated food pellets in adult female rats to model chronic tamoxifen exposure in a clinically-relevant way. Gonadally-intact adult female Long-Evans Hooded rats consumed tamoxifen medicated food pellets for approximately 12 weeks while control animals received standard chow. At the conclusion of the experiment, animals were euthanized, and blood and brain samples were collected for analyses. Blood tamoxifen levels were measured using a novel ultra-performance liquid chromatography-tandem mass spectrometry assay, which found that this administration paradigm produced serum levels of tamoxifen similar to those in human patients. In the brain, brain derived neurotrophic factor (BDNF) was visualized in the hippocampus using immunohistochemistry and quantified using background-subtracted optical densitometry. Chronic oral tamoxifen treatment resulted in a decrease in BDNF expression across several regions of the hippocampus. Together, these findings provide a novel method of modeling and measuring chronic oral tamoxifen exposure, and suggest a putative mechanism by which tamoxifen may cause cognitive and behavioral changes reported by patients.

## Introduction

Tamoxifen is a common adjuvant drug therapy used to decrease the recurrence risk of estrogen receptor-positive breast cancers (Jebahi et al., 2021; Kadakia & Henry, 2015; Novick et al., 2020). The mechanism of action of tamoxifen depends on its location in body tissue and the local estradiol environment (Novick et al., 2020). Generally, tamoxifen acts as an antagonist in the presence of endogenous estradiol, but is agonistic and protects against the effects of estrogen depletion in the absence of endogenous estradiol (Azizi-Malekabadi et al., 2015; Novick et al., 2020). Thus, it has been termed a selective estrogen receptor modulator (SERM). Patients typically take tamoxifen orally for 5-10 years following or concurrent with primary cancer treatment (Kadakia & Henry, 2015).

While tamoxifen is effective at decreasing the risk of cancer recurrence due to its antagonistic effects in breast tissue (Kadakia & Henry, 2015), people undergoing chronic tamoxifen treatment report unwanted psychological side effects such as anxiety, depression, brain fog, confusion, and memory impairment (Jebahi et al., 2021; Lee et al., 2016; Novick et al., 2020; Underwood et al., 2018). Long-term tamoxifen therapy has been shown to decrease performance on visuospatial, visual, and verbal memory tasks (Boele et al., 2015; Castellon et al., 2004; Lejbak et al., 2010; Underwood et al., 2018), decision making tasks (X. Chen et al., 2014; Lejbak et al., 2010), and overall executive function (Boele et al., 2015; Jenkins et al., 2004). However, it has been difficult to separate the effects of tamoxifen from the effects of primary cancer treatments, other adjuvant therapies such as aromatase inhibitors (Breckenridge et al., 2012) and the psychosocial effects of being a cancer survivor (Ahles et al., 2012).

Like in people, tamoxifen treatment in rodents can lead to affective and cognitive changes; however, the impact of tamoxifen on cognitive-behavioral outcomes varies depending on the hormone status of the subjects (Novick et al., 2020). For example, tamoxifen administration to gonad-intact female rodents has been shown to increase anxiety- and depressive-like behaviors (Azizi-Malekabadi et al., 2015), decrease memory consolidation and retrieval ability on step-down avoidance tasks (D. Chen et al., 2002), and decrease novel object recognition ability (Valvassori et al., 2017). In ovariectomized rodents, on the other hand, tamoxifen administration is protective against anxiety- and depressive-like behaviors (Azizi-Malekabadi et al., 2015) and improves working memory ability (Velázquez-Zamora et al., 2012). Because tamoxifen is more commonly used in pre-menopausal patients (Kadakia & Henry, 2015), its effects on gonadally-intact female rodents are of particular interest. Notably, previous rodent studies have not modeled chronic oral tamoxifen use and instead have used a variety of tamoxifen administration methods including long-term injections (Cechinel-Recco et al., 2012), short-term or one-time injections (D. Chen et al., 2002; Kight & McCarthy, 2017; Velázquez-Zamora et al., 2012), and short-term gavages (Valvassori et al., 2017).

The hippocampus is a strong putative candidate to mediate some of the cognitive side effects of tamoxifen therapy. First, the hippocampus expresses _- and β-estrogen receptors (ER_ and ERβ, respectively) across its various subregions (Frick et al., 2015; Harte-Hargrove et al., 2013; Spencer et al., 2008; Woolley, 1998). Research in rodents has demonstrated that estradiol fluctuations impact hippocampal neurophysiology (Liu et al., 2008; Woolley, 1998), which in turn affects depressive- and anxiety-like behaviors (Green & Galea, 2008; Walf & Frye, 2006) as well as performance on spatial memory tasks (Frick et al., 2015; Spencer et al., 2008). Within hippocampal neurons, Brain-Derived Neurotrophic Factor (BDNF) may play an important role in mediating tamoxifen’s effects on cognition and emotion. Estrogen receptor activation influences BDNF transcription via estrogen response elements (EREs) that bind to the *BDNF* gene (Harte-Hargrove et al., 2013; Scharfman & MacLusky, 2006; Sohrabji & Lewis, 2006; Spencer et al., 2008). Indeed, fluctuating levels of estradiol during the estrous cycle lead to fluctuating levels of BDNF expression in the hippocampus (Harte-Hargrove et al., 2013; Luine & Frankfurt, 2013; Scharfman & MacLusky, 2014). Finally, estradiol has been shown to mediate both synaptogenesis (Sato et al., 2007) and LTP (Spencer et al., 2008) through its interactions with BDNF.

Taken together, this evidence suggests tamoxifen’s impact on cognition and emotion may be mediated by BDNF expression in the hippocampus. We therefore hypothesized that long-term oral tamoxifen administration would decrease BDNF expression in the hippocampus. To test this hypothesis, we established a model of long-term oral tamoxifen administration in adult, intact female Long-Evans rats. Using a novel ultra-performance liquid chromatography-tandem mass spectrometry (LC-MS/MS) detection method, we found that this long-term oral administration produced serum levels of tamoxifen comparable to those from Tamoxifen administration in people (Farrar & Jacobs, 2022). Brain tissue analyses found that tamoxifen-treated animals had significantly lower BDNF expression in the DG, medial CA3, and CA1. These results provide an important first step in characterizing the effect of long-term oral tamoxifen administration on the hippocampus, and may increase our understanding of the cognitive and affective side effects associated with long-term tamoxifen use.

## Methods

### Subjects

For all experiments, adult female Long-Evans Hooded rats were purchased from Charles River Laboratories (Wilmington, Massachusetts) at 3-4 months of age. Animals were housed in pairs in standard cages (17.7 x 9.4 in x 8.26 in; Allentown LLC, Allentown, NJ, USA) containing standard TekFresh cellulose low-dust rat bedding (Teklad 7099; Inotiv, West Lafayette, IN, USA), and a Plexiglas tube (Bio-Serv, Flemington, NJ, USA) and wooden block (Bio-Serv, Flemington, NJ, USA) for enrichment. Subjects were kept on a 12-hour light-dark cycle with *ad libitum* food and water access.

### Tamoxifen Self-Administration

Animals were divided into two experimental groups: tamoxifen (n = 20) and control (n = 16). Animals in the tamoxifen group consumed tamoxifen *ad libitum* for 10-13 weeks via medicated food pellets (Envigo, Indianapolis, IN) that were custom developed to simulate the standard dose of tamoxifen prescribed to premenopausal women diagnosed with breast cancer (20 mg/kg daily) while control animals received an ingredient-matched rodent feed (16% protein, 55% carb, 3.4% fat) that did not contain Tamoxifen (Envigo, Farrar & Jacobs, 2022). The composition of the feed was identical between the two groups except for the addition of tamoxifen, red food coloring, and sucrose to enhance palatability in the medicated food pellets (**Table 1**). Importantly, the base pellet (2016, Teklad Global 16% Protein Rodent diet) to which the tamoxifen and associated ingredients were added, and that was fed ad-lib to control rats, did not contain alfalfa or soybean meal, thus minimizing the occurrence of phytoestrogens in control rats during the time they were housed at Haverford College.

**Table 1.** Formula Composition of Medicated and Non-Medicated Pellets. *for detailed formulation of 2016 Teklad Global 16% Protein Rodent Diet, see Inotiv.com.

|  | **Tamoxifen Diet (g/Kg)** | **Control Diet (g/Kg)** |
| --- | --- | --- |
| **2016, Teklad Global 16% Protein Rodent Diet\*** | 949.75 | 1000 |
| **Sucrose** | 49.96 | 0.00 |
| **Tamoxifen USP** | .04 | 0.00 |
| **Red Food Color** | 0.25 | 0.00 |

Body weight was monitored for all animals, regardless of experimental group, throughout the drug administration period. Rats that lost more than 15% of their starting body weight were presented with food pellets mashed in a flavor-enhanced, nutritionally-fortified gel (DietGel Recovery, Clear H20, Portland, ME, USA) in addition to *ad libitum* access to chow to further enhance palatability and encourage ingestion until weight stabilized. Starting at six weeks into the experiment, all rats in the tamoxifen group were presented with this mash daily to maintain adequate health, while also ensuring adequate consumption of Tamoxifen. As an additional check on animal health, starting at 6 weeks, body condition was also assessed daily with the Body Condition Scale (see **Supplementary Figure 1**). All rats in the Tamoxifen group were assigned scores of 3 (well-conditioned) or above. Final weight checks were taken prior to euthanization.

### Blood Analysis

Following 10-13 weeks of access to medicated or control pellets, animals were euthanized via intracardial perfusion (see below). Blood samples were collected from the inferior vena cava immediately prior to perfusion and stored in heparin-coated vacutainer blood collection tubes (BD Medical, Franklin Lakes, NJ, USA) on ice until centrifugation. Samples were centrifuged at 2300 g at 4°C for 20 min and serum was stored at -80_ until assay.

An ultra-performance liquid chromatography-tandem mass spectrometry assay with reversed-phase chromatographic separation employing a Waters XBridge C18 (100 x 2.1 mm, 100 Å, 3.5 µm) and a runtime of 4.5 minutes was used to quantify tamoxifen levels. Rat plasma samples (100 µL) were extracted with acetonitrile containing tamoxifen-d5 (5 ng/mL) and 0.1% formic acid. Two µL of the extract were injected onto the UPLC-MS/MS system for analysis.

Tamoxifen and tamoxifen-d5 were separated by ultra-performance liquid chromatography (UPLC) and detected using a triple quadrupole mass spectrometer (API4000). Tandem mass spectrometery (MS-MS) parameters were optimized in the positive ionization mode and multiple reaction monitoring (MRM) transition of m/z 372.3 72.0 was used for tamoxifen analysis. Tamoxifen-d5 (m/z 377.3 ➔ 72.0) was used as an internal standard. A calibration curve for tamoxifen was prepared in human plasma over the linear range of 0.1 ➔ 250 ng/mL (r^2^ > 0.99, Figure 2, Table 2). Calibration standards and blanks in human plasma were extracted by protein precipitation with 400 μL of 5 ng/mL of tamoxifen-d5 in acetonitrile. Five μL of extract was injected for LC-MS/MS analysis. Tamoxifen and tamoxifen-d5 were separated on an XBridge C18 column (100 x 2.1 mm, 100 Å, 3.5 μm) using 0.1 % formic acid in water and 0.1 % formic acid in acetonitrile as aqueous and organic mobile phases with a flow rate of 0.5 mL/min. Analytes were separated using a gradient elution, with retention time of 2.68 min for tamoxifen and tamoxifen-d5. The LC-MS Tamoxifen assay used here was based on assays initially developed and validated in human plasma (Bobin-Dubigeon et al., 2019; Rama Raju et al., 2015). Therefore, as a confirmatory assay, human plasma and rat plasma were compared using 100 ng/mL concentrations of tamoxifen and found to be comparable (**Table 3**).

**Figure 1.**
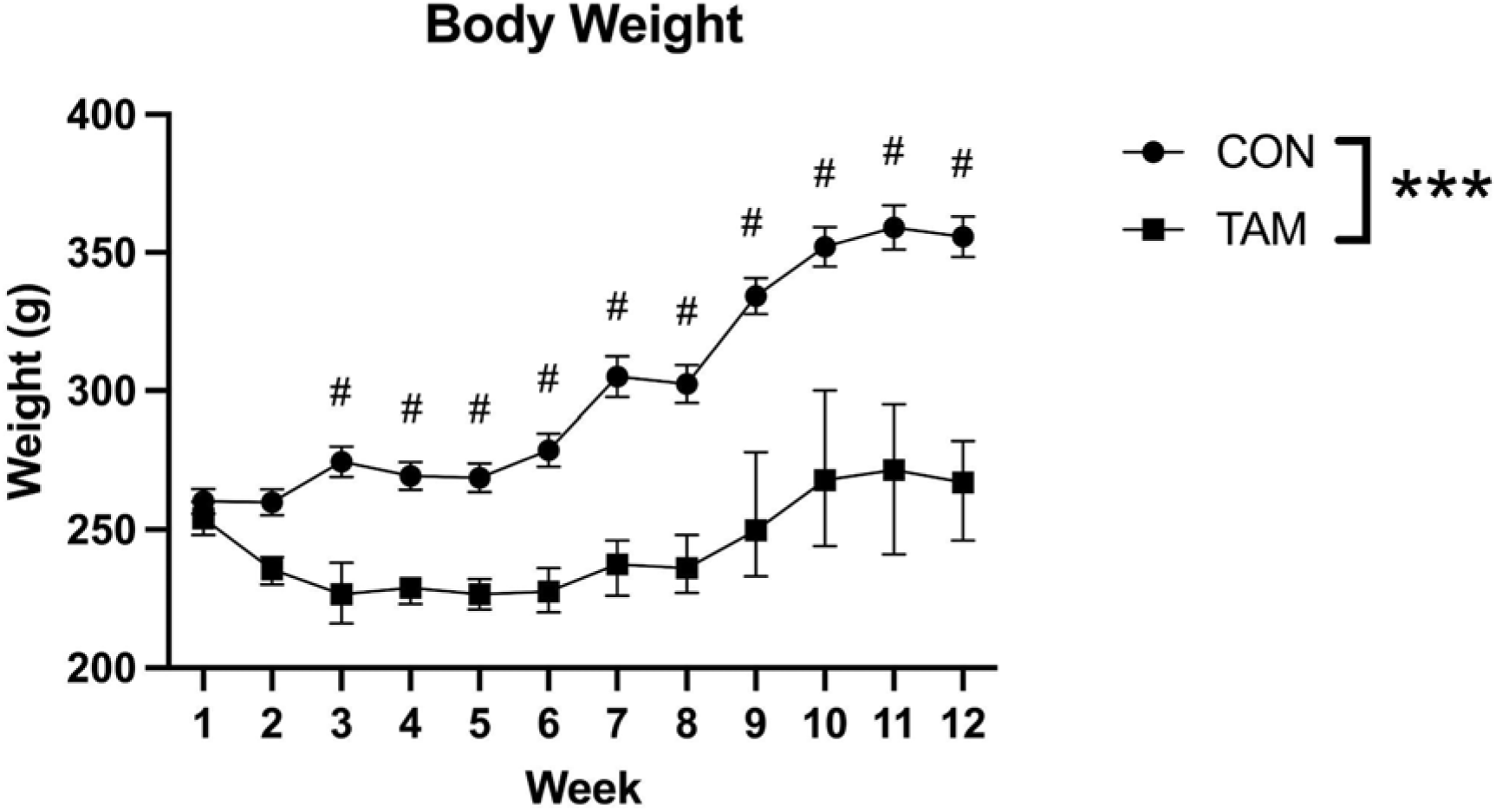
Body Weight. Body weight was tracked throughout the course of the experiment. Starting at Week 6, Tamoxifen pellets were mashed in a flavor-enhanced gel in addition to *ad libitum* presentation to prevent further weight loss. Repeated measures factorial ANOVAs were used to look for interactions and main effects of drug condition (TAM vs. CON) and time (Week) on body weight. There was a significant interaction between drug condition and time on body weight. Post-hoc tests found that starting at Week 3, TAM animals weighed significantly more than CON animals. Data presented as mean ± standard error, ***significant interaction, p < 0.001; # significant post hoc test, p < 0.001.

**Figure 2.**
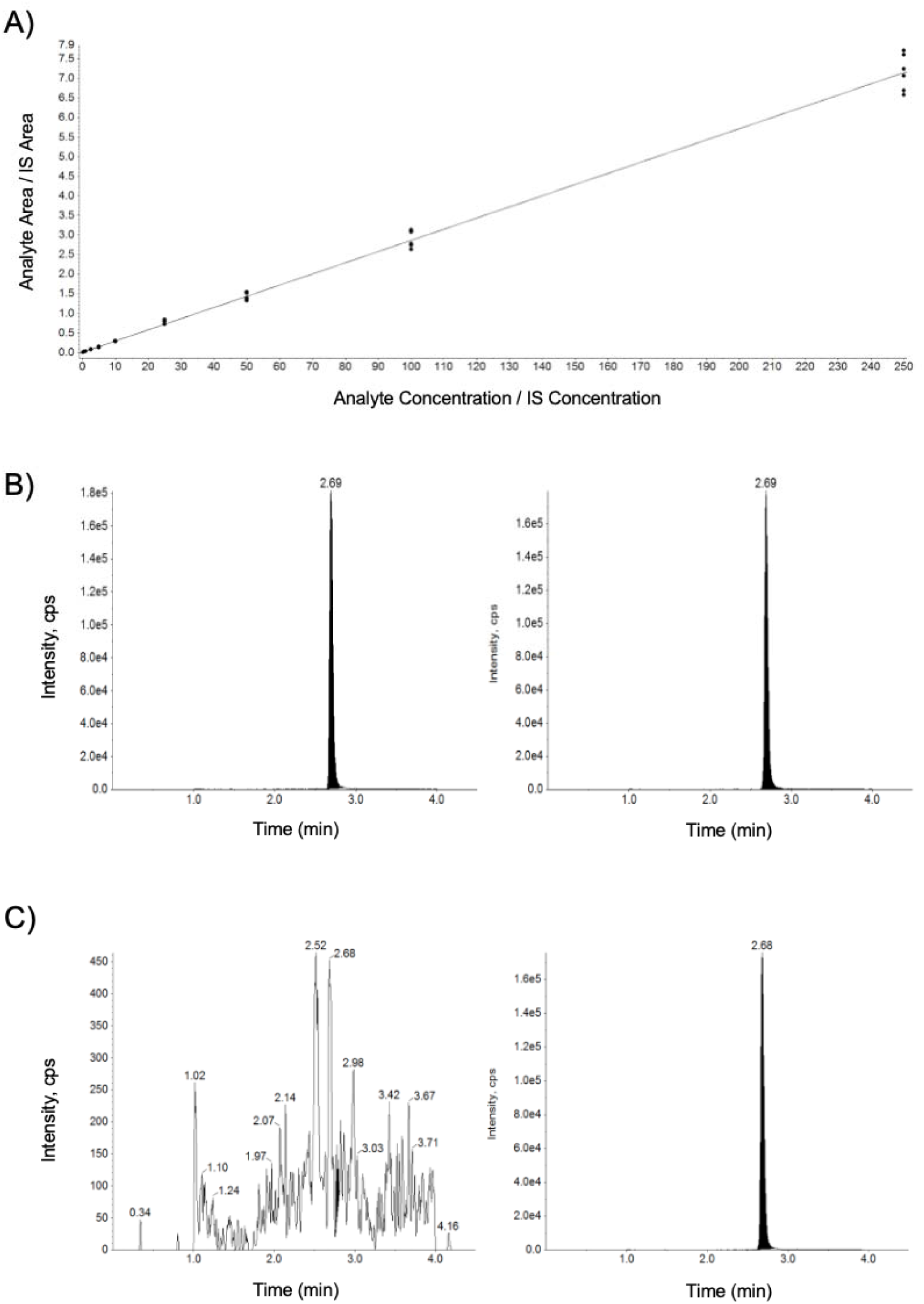
Mass Spectrometry Analysis of Tamoxifen. Tamoxifen and tamoxifen-d5 were separated by ultra-performance liquid chromatography (UPLC) and detected using a triple quadrupole mass spectrometer. Tandem mass spectrometry (MS-MS) parameters were optimized in the positive ionization mode and the multiple reaction monitoring (MRM) transition of m/z 372.3 72.0 was used for tamoxifen analysis. Tamoxifen-d5 (m/z 377.3 72.0) was used as an internal standard. A) A calibration curve for tamoxifen was prepared in human plasma and was linear over the range of 0.1 – 250 ng/mL. Analytes were separated using a gradient elution, with retention time of 2.68 min for tamoxifen and tamoxifen-d5. B) Representative chromatogram from Tamoxifen-treated animal of Tamoxifen (left) and Tamoxifen-d5 (right). C) Representative chromatogram from control-treated rat (standard chow) of Tamoxifen (left) and Tamoxifen-d5 (right) in plasma.

**Table 2.** Summary of Tamoxifen calibration curve in plasma. A calibration curve for tamoxifen was prepared in human plasma and was linear over the range of 0.1 – 250 ng/mL with coefficient of regression, r^2^ > 0.99. Data presented as Mean, Standard Deviation (SD), Accuracy (%), and Coefficient of Variation (CV %).

| Calibration Curve (n=6) | Tamoxifen Concentration (ng/mL) |  |  |  |  |  |  |  |  |  |
| --- | --- | --- | --- | --- | --- | --- | --- | --- | --- | --- |
|  | 0.1 | 0.5 | 1 | 2.5 | 5.0 | 10 | 25 | 50 | 100 | 250 |
| 1 | 0.108 | 0.538 | 1.00 | 2.61 | 5.11 | 10.5 | 29.4 | 46.4 | 108 | 270 |
| 2 | 0.117 | 0.533 | 1.11 | 2.61 | 4.84 | 10.3 | 27.4 | 48.4 | 107 | 253 |
| 3 | 0.104 | 0.484 | 0.95 | 2.69 | 4.79 | 10.2 | 29.2 | 54.0 | 109 | 247 |
| 4 | 0.090 | 0.479 | 0.88 | 2.08 | 4.25 | 9.55 | 25.2 | 47.9 | 95.6 | 266 |
| 5 | 0.091 | 0.566 | 0.922 | 2.49 | 5.06 | 10.5 | 27.9 | 52.9 | 92.1 | 234 |
| 6 | 0.092 | 0.467 | 0.841 | 2.18 | 4.25 | 9.38 | 24.8 | 46.5 | 97.2 | 230 |
| Mean | **0.100** | **0.511** | **0.949** | **2.44** | **4.72** | **10.1** | **27.3** | **49.4** | **101** | **250** |
| SD | 0.011 | 0.040 | 0.096 | 0.253 | 0.382 | 0.487 | 1.95 | 3.29 | 7.35 | 16.3 |
| Accuracy (%) | **100** | **102** | **94.9** | **97.7** | **94.3** | **101** | **109** | **98.7** | **101** | **100** |
| CV (%) | 11.1 | 7.79 | 10.1 | 10.4 | 8.09 | 4.84 | 7.15 | 6.66 | 7.25 | 6.52 |

**Table 3.**
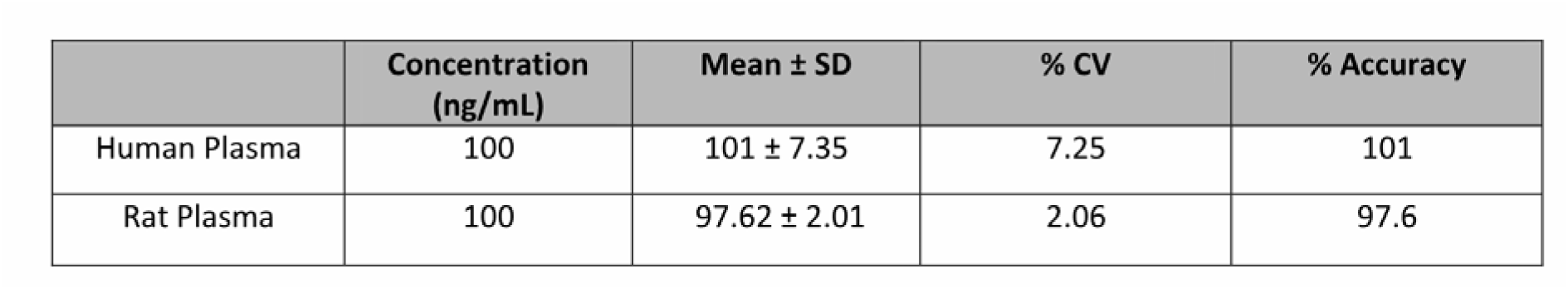
Human plasma and rat plasma comparison. The Tamoxifen assay was initially developed in human plasma. Human plasma and rat plasma were compared using 100 ng/mL concentrations of tamoxifen. Results were comparable between human and rat plasma. Concentration (ng/mL), Data presented as Concentration (ng/mL), Mean, Standard Deviation (SD), Coefficient of Variation (% CV), and Accuracy (%).

| | Concentration<br>(ng/mL) | Mean ± SD | % CV | % Accuracy |
| --- | --- | --- | --- | --- |
| Human Plasma | 100 | 101 ± 7.35 | 7.25 | 101 |
| Rat Plasma | 100 | 97.62 ± 2.01 | 2.06 | 97.6 |

### Tissue Histology and Immunohistochemistry

At the conclusion of the experiment, subjects were sacrificed by intracardial perfusion. Briefly, subjects were given an overdose of sodium pentobarbital (Beuthanasia-D, 22 mg/100 g body weight, Merck Animal Health, Madison, NJ, USA) and then transcardially perfused with approximately 250 mL of 25mM phosphate buffered saline (PBS) followed by approximately 250 mL of 4% paraformaldehyde in PBS. Brains were immediately extracted and post-fixed in 4% paraformaldehyde overnight and then cryoprotected for 48 h in 30% sucrose in PBS. Coronal sections (35 µm) were sectioned at -20°C on a cryostat (Leica Biosystems, Wetzlar, Germany), collected in a 1:6 series, and stored in cryoprotectant until immunohistochemical processing.

For immunohistochemistry, hippocampal sections were removed from cryoprotectant and rinsed 5 x 5 minutes in 25 mM PBS. After PBS washes, sections were incubated in hydrogen peroxide (1:100) for 15 minutes to reduce endogenous peroxidase activity. Sections were then washed 5 x 5 minutes in 25 mM PBS and before incubation in a rabbit monoclonal primary antibody against brain-derived neurotrophic factor (1:3000, Cat # ANT-010, Alomone Labs, Jerusalem, Israel) in 0.4% Triton-X 100 in PBS for 24 hours at room temperature. This antibody has previously been validated for use in rodent tissue using inducible BDNF knockouts (Sun et al., 2018) and shBDNF treated rats (Taliaz et al., 2010). This antibody recognizes both mature BNDF and pro-BDNF (Taliaz et al., 2010).

After incubation in the primary antibody, sections were rinsed in PBS (5 x 5 minutes) and incubated for one hour in a biotinylated secondary antibody (1:600, Biotin-SP AffiniPure Goat Anti-Rabbit IgG, Jackson Immunoresearch Labs; 111-065-003) in 0.4% Triton-X 100 in PBS. Sections were then rinsed in PBS (5 x 5 minutes) and incubated in avidin-biotin complex with horseradish peroxidase (per manufacturer’s instructions, Vectastain Elite ABC-HRP Kit, Vector Laboratories, Burlingame, CA, USA). After 4 x 5 minute washes in PBS, sections were washed 2 x 5 minutes in a sodium acetate buffer and then incubated for 10 minutes in 3,3’-diaminobenzidine HCl (0.2 mg/mL) in a nickel sodium acetate buffer (0.025 g Nickel II Sulfate/mL NaOAc) and hydrogen peroxide (0.83 µL/mL) solution.

Stained tissue was then washed in sodium acetate buffer (2 x 5 min), washed in PBS (3 x 5 min) mounted on glass slides (SuperFrost Plus Microscope Slides, ThermoFisher Scientific, Waltham, MA, USA) and left to air-dry overnight. After drying, slides were dehydrated with alcohols, cleared with xylenes and coverslipped with Permount (ThermoFisher Scientific, Waltham, MA, USA).

### BDNF Densitometry Analysis in Hippocampus

Images were acquired at 10X magnification using a Nikon E200 brightfield microscope with a color camera and Spot Basic software (Diagnostic Instruments, Sterling Heights, MI, USA). Five evenly-spaced template images (S1, S2, S3, S4, S5) were selected from the rat brain atlas (Paxinos & Watson, 2013) to capture the rostral to caudal extent of the hippocampus. The CA3 was split into medial and lateral regions for imaging and analysis in order to capture it fully. Twenty images were collected for each animal: five images per hippocampus region (DG, CA1, mCA3, lCA3).

Densitometry analyses were performed by experimenters who were blind to the condition of the subject using ImageJ software (imagej.net/software/fiji; Schindelin et al., 2012). Mean values for background optical density were collected from 36×36 pixel regions that did not contain BDNF-ir cells using the “mean gray value” tool. Mean optical density values for regions of interest (ROIs) were calculated after tracing the hippocampus region of interest with the polygon tool (**Figure 3**) and analyzing with the “mean gray value” tool (**Figure 4**). ROIs were checked for consistency across animals by an experimenter blind to the condition of the subjects. Background optical density was subtracted from ROI optical density using the following equation: (OD) = log (255 / (OD_ROI_ - OD_background_)). Because background levels were variable across sections, likely due to individual differences in perfusions, we background-subtracted OD values between sections with high background levels versus low background levels and found no significant difference between groups. This suggests that: 1) our method was effective at subtracting background and 2) differences between treatment groups were not due to differences in background.

**Figure 3.**
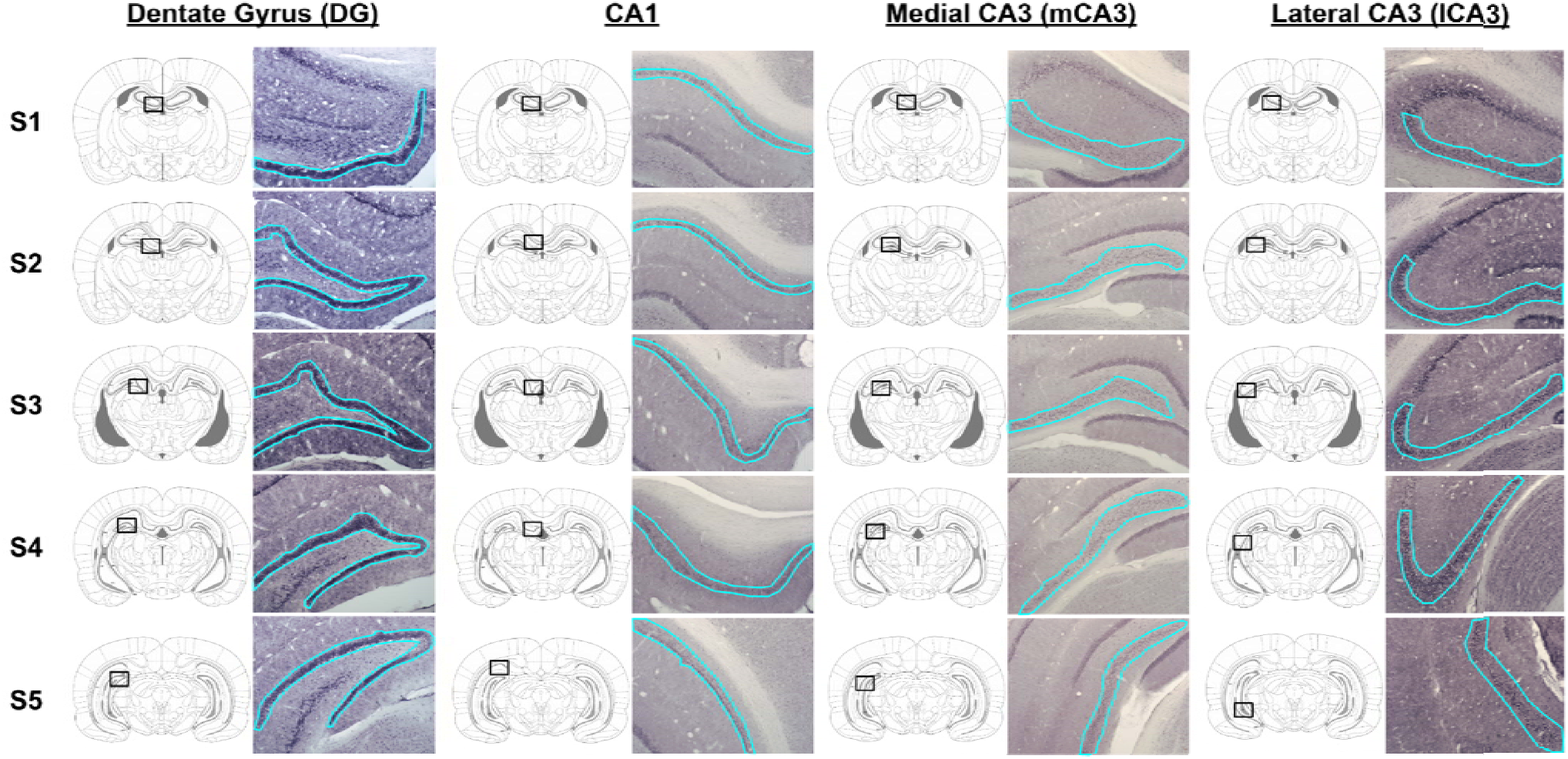
BDNF Expression in Hippocampal Subregions. Representative photomicrographs showing immunohistochemical localization of BDNF in the dentate gyrus (DG), CA1, medial CA3 (mCA3) and lateral CA3 (lCA3). For each region of interest, five sections (S1-S5) capturing the rostral to caudal extent of the hippocampus were analyzed. Regions of interest are identified on the corresponding template atlas plate and traced in cyan on the photomicrograph.

**Figure 4.**
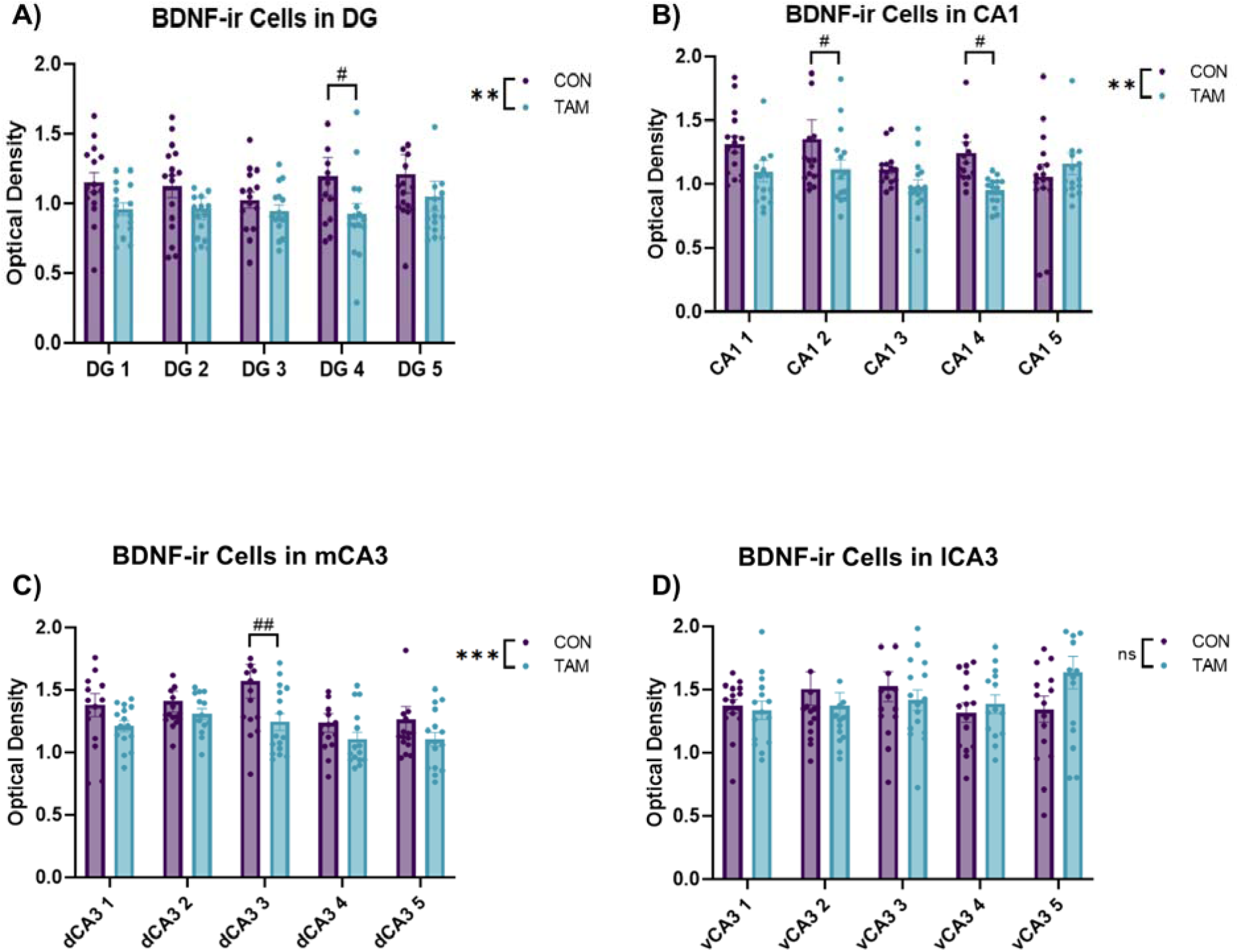
Quantification of BDNF Optical Density in Hippocampal Subregions. For each brain region of interest, immunohistochemical localization of BDNF was quantified using background-subtracted optical density in five sections capturing the rostral to caudal extent of the hippocampus. Significant interactions and/or main effects of drug condition (Tamoxifen, TAM vs. Control, CON) and section number (1-5) on BDNF optical density were detected using two-way ANOVAs. A) In DG, there was a significant main effect of drug condition on BDNF-ir density. Post-hoc tests revealed that BDNF-ir density was lower in TAM animals than CON animals in Section 4. B) In CA1, there was a main effect of drug condition on the density of BDNF-ir cells. Post hoc tests revealed that BDNF-ir density was lower in TAM animals than CON animals in section 2. C) In mCA3, there was also a significant main effect of drug condition on the density of BDNF-ir cells. Post-hoc tests revealed that BDNF-ir was lower in TAM animals than CON animals in Section 3. D) Finally, in lCA3, there were no interactions or main effects of drug condition or section level on BDNF immunoreactivity. Data are presented as mean ± standard error, asterisks indicate significant omnibus tests, * p < 0.05; ** p < 0.01; *** p < 0.001. Pound signs indicate significant post-hoc differences between drug conditions, # p < 0.05; ## p < 0.01; ### p < 0.001

## Statistical Analysis

All statistics were run using Jamovi (www.jamovi.org) with alpha values of 0.05. Assumptions of normality were tested using the Shapiro-Wilk normality test. Repeated measures factorial ANOVAs were used to look for interactions and main effects of drug condition (tamoxifen vs. control) and time (week) on body weight. Significant effects were explored using Bonferroni’s multiple comparisons tests and the Geisser and Greenhouse method was used to correct violations of sphericity. Two-way ANOVAs were used to look for interactions and main effects of drug condition (tamoxifen vs. control) and section level (1-5) on BDNF-ir density within each of the four ROIs (DG, CA1, vCA1, and CA3). Individual brain regions were analyzed separately. Significant effects were explicated using Fisher’s LSD post-hoc comparisons. Individual samples were excluded from analysis if tissue damage precluded OD analysis, or if there was no adequate match for the template atlas plate. The relationship between serum tamoxifen levels and BDNF-ir density was examined using simple correlations.

## Results

### Body Weight

There was a significant interaction between time and drug condition on body weight (*F*(11,99 = 10.65, p < 0.0001). Post-hoc tests revealed that starting at Week 3, Tamoxifen-fed rats weighed significantly more than control rats for the duration of the experiment (all p < 0.001, **Figure 1**).

### Plasma Tamoxifen Levels

Tamoxifen was not found in any control samples. In tamoxifen-treated animals, plasma concentrations ranged from 10.7 to 48.3 ng/mL (M = 28.1, SD = 11.2, Table 4). Representative chromatograms can be found in Figure 2.

**Table 4.**
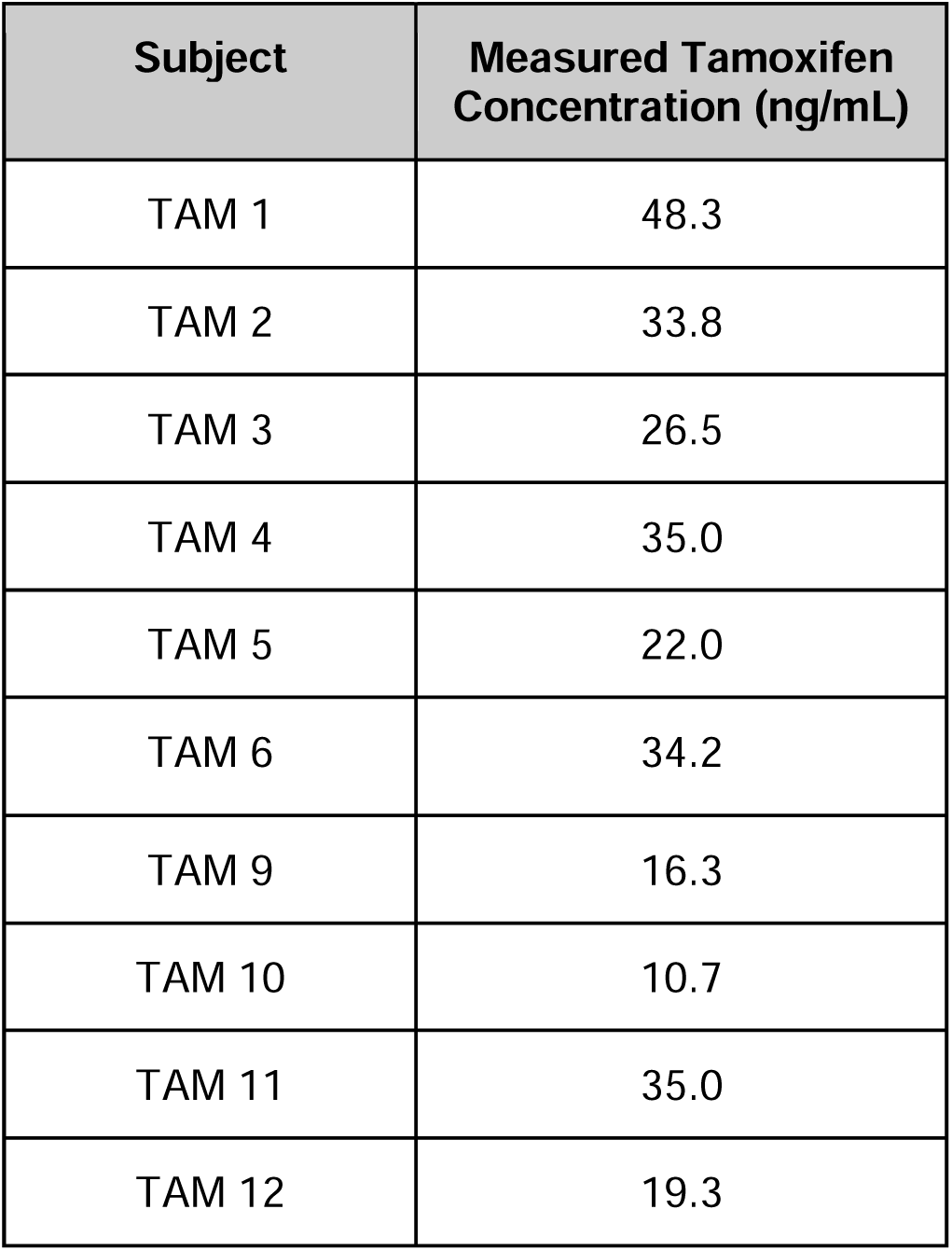
Plasma Tamoxifen Concentration. Tamoxifen levels (ng/mL) in TAM-treated rats (*n* = 10).

| Subject | Measured Tamoxifen Concentration (ng/mL) |
| --- | --- |
| TAM 1 | 48.3 |
| TAM 2 | 33.8 |
| TAM 3 | 26.5 |
| TAM 4 | 35.0 |
| TAM 5 | 22.0 |
| TAM 6 | 34.2 |
| TAM 9 | 16.3 |
| TAM 10 | 10.7 |
| TAM 11 | 35.0 |
| TAM 12 | 19.3 |

### BDNF Immunoreactivity

For each brain region of interest, the density of BDNF-immunoreactive (ir) neurons in the DG, mCA3, lCA3, and CA1 was analyzed and compared between groups. In the DG, there was a significant main effect of drug condition on the density of BDNF-ir cells (F(1, 146) = 9.454, p = 0.003). Post-hoc tests revealed that BDNF-ir density was lower in tamoxifen-treated animals than control animals in Section 4 (p = 0.03, **Figure 4A**). In CA1, there was a main effect of drug condition on the density of BDNF-ir cells (F(1, 146) = 8.886, p = 0.003). Post hoc tests revealed that BDNF-ir density was lower in tamoxifen-treated animals than control animals in section 2 (p = 0.04) and section 4 (p = 0.02, **Figure 4B**). In mCA3, there was a significant main effect of drug condition on the density of BDNF-ir cells (F, 1, 144) = 11.94, p < 0.001). Post-hoc tests revealed that BDNF-ir was lower in tamoxifen-treated animals than control animals in Section 3 (p < 0.01, **Figure 4C**). There was also a significant main effect of section on the density of BDNF-ir cells in mCA3 (F(4, 144) = 3.302, p = 0.01). Post-hoc tests revealed that Section 2 differed from Section 4 (p = 0.02) and Section 5 (p = 0.03), and Section 3 differed from Section 4 (p < 0.01) and Section 5 (p < 0.01). Finally, in lCA3, there was no effect of drug condition (F(1, 145) = 0.071, p = 0.79) or section (F(1,145) = 0.87, p = 0.48) on the density of BDNF-ir cells (**Figure 4D**).

Animals’ plasma tamoxifen concentration at the time of sacrifice did not correlate significantly with BDNF-ir cells in the DG (r = -.4221, p = .2244, **Figure 5A**) or CA1 (r = - .2053, p = .5693, **Figure 5B**). In both medial and lateral CA3 (**Figure 5C and 5D**), there was a trend towards increased plasma tamoxifen concentration being associated with decreased BDNF-ir cells, although these did not reach significance (mCA3 r = -.6112, p = .0604; lCA3 r = -.6168, p = .0575).

**Figure 5.**
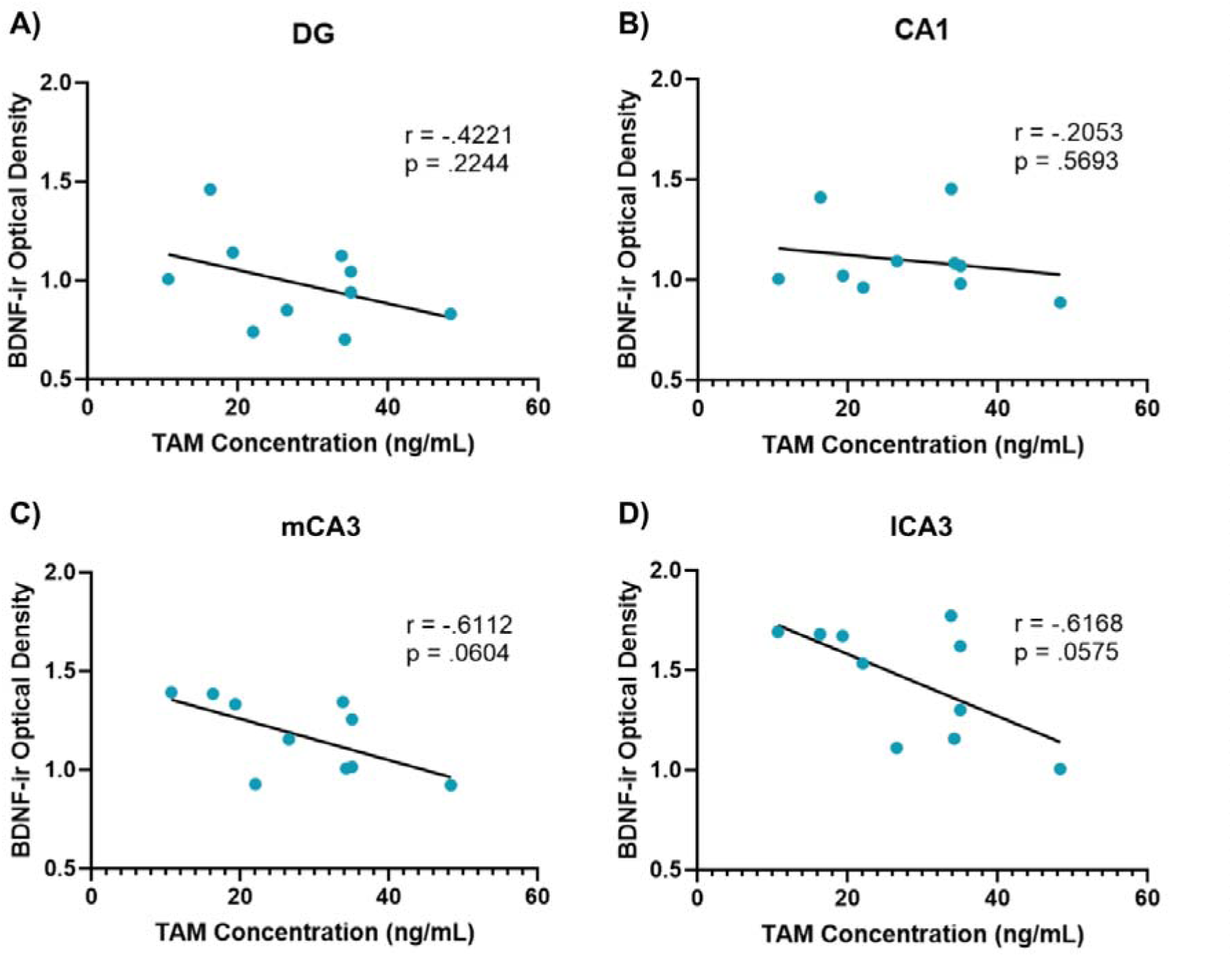
Plasma Tamoxifen and Brain BDNF Correlations. The relationship between plasma tamoxifen levels and BDNF expression in hippocampal subregions was investigated using simple correlations. In all four brain regions of interest, plasma tamoxifen was negatively correlated with brain BDNF levels. Although none of these correlations reached significance, trends towards significance were detected in C) mCA3 and D) lCA3. Pearson’s correlation coefficient (r) and p values for each brain area presented.

## Discussion

Here we demonstrate for the first time that long-term oral tamoxifen administration decreases BDNF expression in the hippocampus of adult female Long-Evans Hooded rats. Notably, we show that a novel administration method, long-term *ad lib* exposure to medicated food pellets, produces plasma tamoxifen levels that are similar to a human dose (20 mg/kg; Farrar & Jacobs, 2022). Further, we used a novel LC-MS/MS assay to detect tamoxifen levels in rat plasma. Together, these results provide a new way to model and measure the effects of long-term oral tamoxifen administration on the brain, which may be particularly useful for investigating mechanisms of cognitive and behavioral changes reported by patients taking Tamoxifen.

Voluntary oral administration via medicated food pellets is a clinically relevant, non-invasive, and effective method for modeling the effects of tamoxifen in rodents. Tamoxifen can readily cross the blood-brain-barrier (Lien et al., 1991; Pareto et al., 2004), and oral self-administration is an attractive alternative to systemic injections (Cechinel-Recco et al., 2012; D. Chen et al., 2002; Kight & McCarthy, 2017; Velázquez-Zamora et al., 2012) or oral gavage (Valvassori et al., 2017) in that it more closely models the metabolism of tamoxifen use in patient populations and eliminates stress associated with restraint, repeated injections, or gavage as potential confounding variables. Nonetheless, there are some challenges with this method. First, there were likely differences in palatability between tamoxifen and control chow. Tamoxifen-fed animals initially lost weight and required additional intervention (food pellets mashed in flavor-enhanced gel) to prevent further weight loss, whereas control animals gained weight across the course of the experiment. A similar study in mice found that 15 days of exposure to tamoxifen chow also led to weight loss, particularly in female subjects, and animals did not consistently eat Tamoxifen chow for the first 5 days of exposure (Smith et al., 2022). However, an anorexic effect of tamoxifen itself cannot be ruled out, as tamoxifen injections induce anorexia in rodents (Lampert et al., 2013; Gray et al., 1993; López et al., 2006; Wade & Heller, 1993; Walker et al., 2011, p. 2006). While oral gavage would introduce stress and is therefore not recommended, future experiments may wish to explore mild food deprivation, pair feeding tamoxifen-treated and control animals, or giving a daily oral dose of tamoxifen to eliminate the effects of feeding differences and/or body weight as potential confounds on dependent variables. There was also variability in individual animals’ plasma tamoxifen concentrations. These differences likely resulted from differences in voluntary consumption of tamoxifen chow across animals, both throughout the experiment and specifically on the day of euthanasia. Blood draws were collected approximately 60 minutes after the last tamoxifen pellets were presented, but the amount of medicated pellet consumed, or the timing of consumption within the one-hour window, were not measured. Future studies should collect data about volume and timing of food intake, particularly prior to blood collection, as well as additional bioassays that correlate with tamoxifen consumption such as uterine weight (Fong et al., 2007).

Our finding that long-term oral tamoxifen administration decreased BDNF expression in the hippocampus may point towards a putative mechanism underlying cognitive side effects that patients taking tamoxifen report. Given the known association of estrogen receptor activation and BDNF transcription (Harte-Hargrove et al., 2013; Scharfman & MacLusky, 2006; Sohrabji & Lewis, 2006), it is possible that tamoxifen’s action at estrogen receptors in the brain could disrupt BDNF transcription, altering plasticity and neuronal stability in the hippocampus. These changes to hippocampal function could in turn impair cognitive function and lead to clinical symptoms like brain fog, memory deficits, and alterations in mood/anxiety. In support of this idea, Smith et al. (2022) found that short term Tamoxifen administration impacts hippocampal neurogenesis. Specifically Tamoxifen injection is associated with decreased progenitor cell proliferation in the dentate gyrus, although Tamoxifen ingestion is associated with increased neuronal differentiation in the dentate gyrus. Thus, our finding that long-term oral tamoxifen administration decreased BDNF density in the current study may be due to a decrease in cell proliferation, or conversely, decreases in BDNF following Tamoxifen treatment could contribute to changes in neurogenesis.

Brain BDNF is difficult to visualize due to its relatively low expression in mature neurons. In the current study, we used an immunohistochemistry approach with a nickel-enhanced chromogen reaction to amplify visualization. This method produces robust BDNF staining, but can produce higher levels of background staining, necessitating a background-subtracted densitometry approach to quantification rather than individual cell counts. Further, BDNF staining patterns differ across published experiments, likely due to differences in antibodies. Here, we report BDNF expression in the granular cell layer is reduced following long-term oral Tamoxifen administration, but found little BDNF expression in the mossy fiber layer. This matches the expression pattern previously shown with this antibody (Taliaz et al., 2010). However, previous papers that used different antibodies report BDNF staining in the mossy fiber layer, but only sparse staining in the granule cell layer (Conner et al., 1997; Dieni et al., 2012). It is possible that different antibodies label distinct pools of BDNF based on differences in the targeted epitope. The method of visualization and antibody choice are therefore important considerations when visualizing and quantifying BDNF in the brain.

Future studies using this model of long-term oral tamoxifen administration should include behavioral assays of cognition, mood, and anxiety. Our finding that long-term tamoxifen administration decreased BDNF in the DG, CA1, and medial CA3 suggests that pattern separation, spatial memory, associative memory, and episodic memory processes could be particularly affected (Bartsch et al., 2011; Cherubini & Miles, 2015; Kesner, 2007; Langston et al., 2010). Disparate findings in CA3 subregions further point towards tamoxifen’s effects on cognitive function. For example, there is some literature to suggest that the dorsal CA3 is responsible for cognition and information processing, while the ventral region is more closely associated with stress and affect (Fanselow & Dong, 2010). A full behavioral profile of animals during long-term oral tamoxifen treatment, such as further assays of cognition (e.g., object-in-place, attentional set shifting), anxiety-like behaviors (e.g., defensive burying, light-dark box) and learned helplessness or anhedonia (e.g., forced swim, tail suspension, splash test, sucrose preference) would be useful to characterize tamoxifen’s effects on behavior in this model, and may suggest additional brain regions for exploration. Further, while this study focused on the hippocampus, it is likely that tamoxifen impacts other estrogen-sensitive brain regions that are associated with cognition, anxiety, and/or mood disorders, such as the nucleus accumbens and amygdala. Tamoxifen could disrupt BDNF transcription in these regions by interacting with estrogen receptors via the same putative mechanism that we have identified in the hippocampus. The relationship between Tamoxifen, hippocampal BDNF levels, and body weight also requires further investigation, as BDNF is known to regulate energy balance via action in other parts of the brain (for review see Xu & Xie, 2016).

In addition to a novel administration paradigm, we also used a novel, sensitive, and reproducible method for the analysis of tamoxifen that is suitable for rat plasma. This will be useful to future researchers interested in the physiological effects of oral tamoxifen administration. While not significant, we did observe a trend towards correlation between plasma tamoxifen concentration and BDNF density in mCA3 and lCA3, supporting tamoxifen’s potential role in decreasing BDNF transcription in the hippocampus. It is worth noting that tamoxifen is also commonly used in neuroscience research as a tool to induce Cre mechanisms. While these research applications are markedly different in that the doses are generally much higher and administration is acute (often a single injected dose) (Donocoff et al., 2020; Heffner, 2011) our research suggests that tamoxifen administration affects BDNF expression, and researchers using tamoxifen in their work should consider its potential off-target effects on neurobiology.

Tamoxifen remains the gold-standard medication for preventing recurrence of estrogen receptor-positive breast cancers. Its ongoing use as an adjuvant therapy will likely continue despite common affective and cognitive side effects; tamoxifen treatments will enable breast cancer survivors to live longer and healthier lives. Further identifying and understanding the neural mechanisms underlying these side effects, however, could lead to promising interventions that would mitigate unwanted psychological changes without risking cancer recurrence. It is important that researchers and clinicians continue to explore ways to improve psychiatric health in at-risk and under-researched patient populations. Cancer survivors, women and other gender minorities in particular, are particularly vulnerable to a range of affective, anxiety, and cognitive disorders (Ahles et al., 2012; Di Giacomo et al., 2016; Fafouti et al., 2010). It is imperative that we find ways to support all aspects of their health as they navigate treatment.

## Acknowledgements

This work was funded in part by Haverford College Department of Psychology. The authors wish to thank Shanice Edwards, Christina Vedar, Abigail Cohen, Athena Zuppa, and Ganesh Moorthy from the Children’s Hospital of Philadelphia Research Institute Bioanalytical Core for developing and conducting the mass spectrometry assays. Some of the mass spectrometry data were previously presented as a poster at the American Society for Mass Spectrometry Conference on Mass Spectrometry and Allied Topics in Philadelphia, PA in November of 2021. The authors also wish to thank Dr. Louise Charkoudian for her assistance interpreting the LC-MS data.

---

**Supplementary Figure 1.**
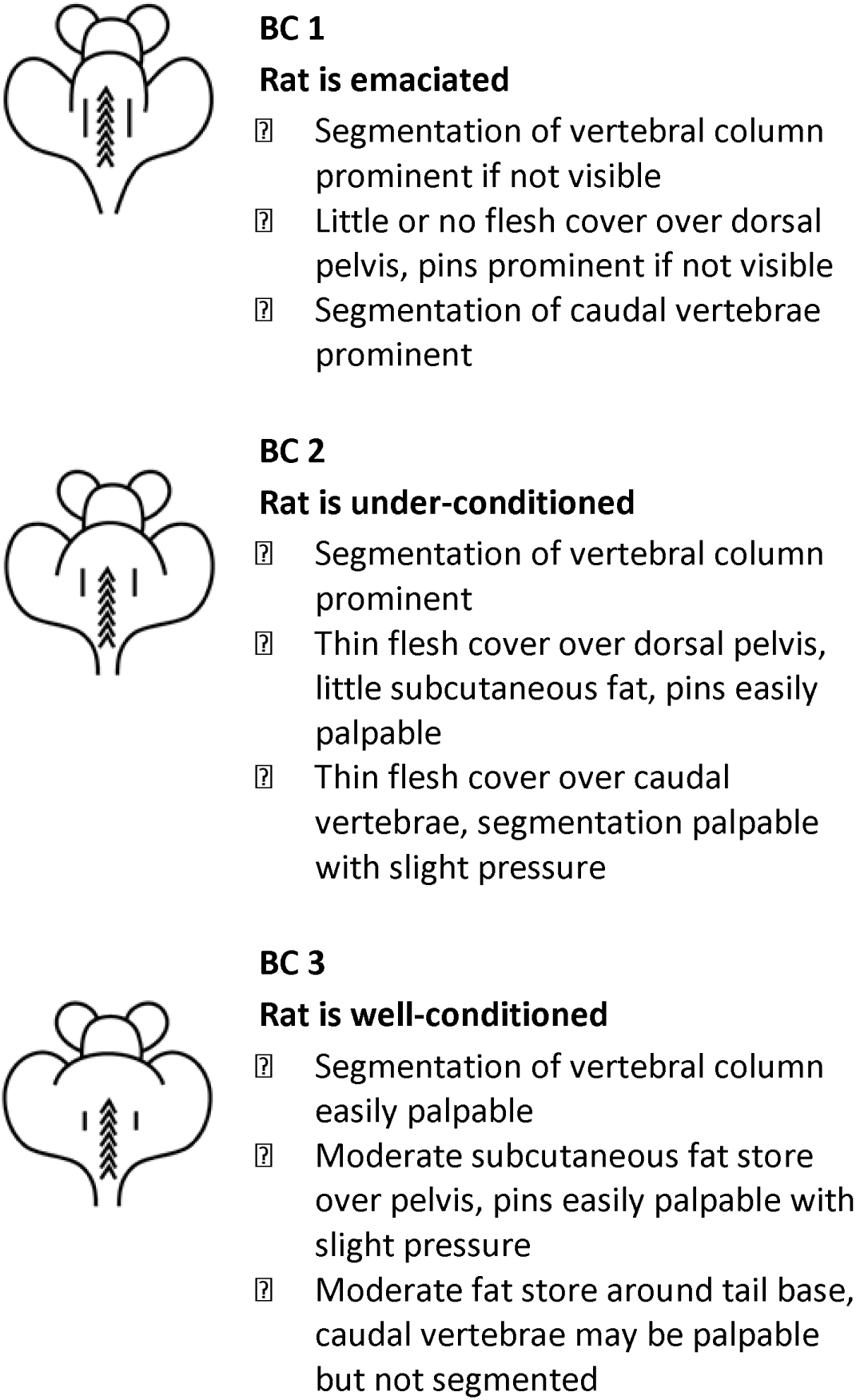
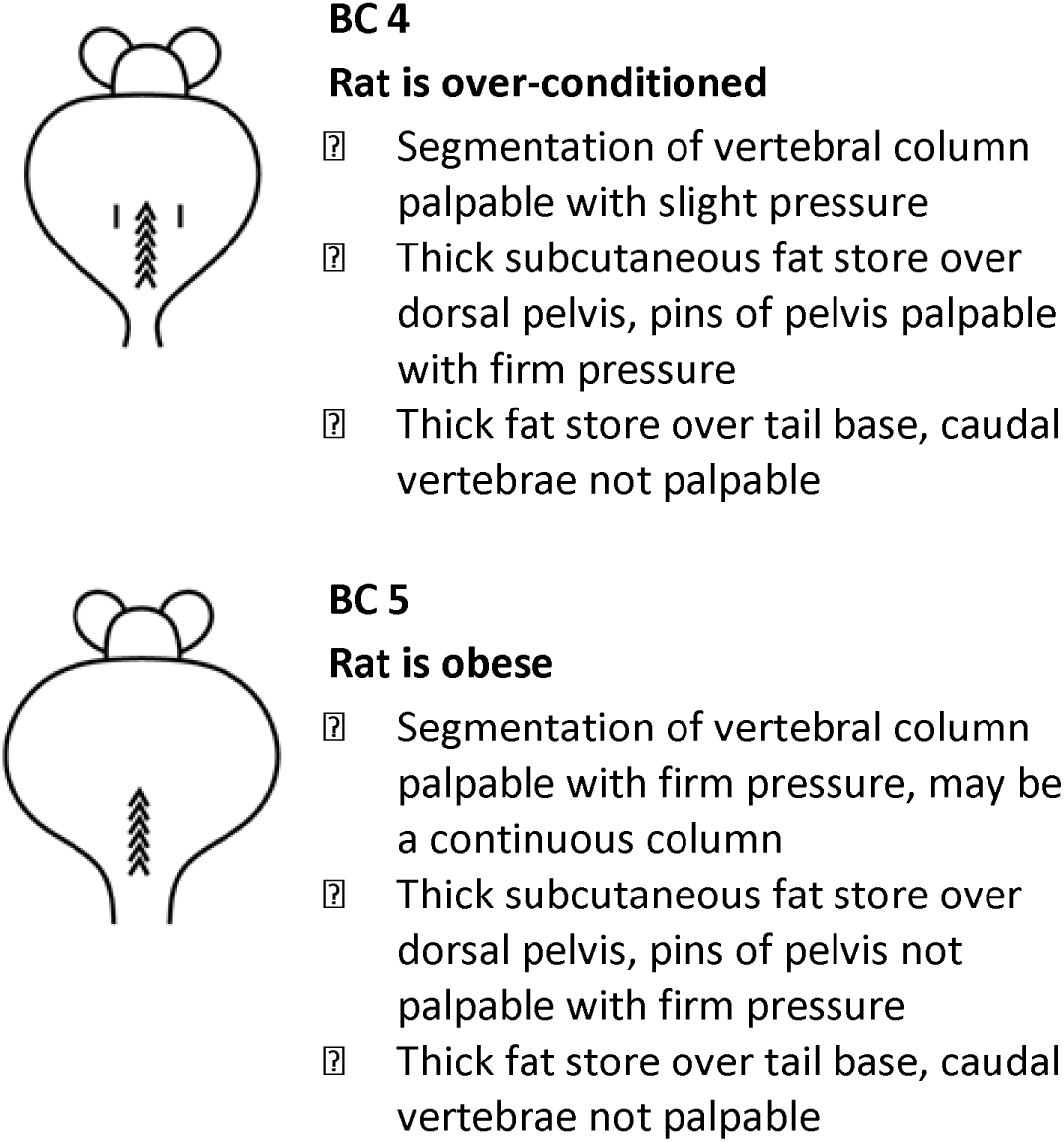
Body Condition Scoring. In addition to weight, Body Condition (BC) Scoring was used as a measure of animal health. Adapted from: Hickman D, Swan M. 2010. Use of a Body Condition Score Technique to Assess Health Status in a Rat Model of Polycystic Kidney Disease.

